# Hyperspectral imaging of Marchantia

**DOI:** 10.64898/2026.05.28.721262

**Authors:** Grace Zi Hao Tan, Daisuke Urano

**Affiliations:** Temasek Life Sciences Laboratory, 1 Research Link, National University of Singapore, Singapore 117604; Disruptive & Sustainable Technologies for Agricultural Precision (DiSTAP) IRG, Singapore-MIT Alliance of Research and Technology, Singapore 138602; Department of Biological Sciences, National University of Singapore, 16 Science Drive 4, Singapore 117558

**Keywords:** Hyperspectral imaging, Marchantia, nutrient deficiency, plant diagnostics, pigmentation patterns

## Abstract

Hyperspectral imaging is an imaging technique that allows for acquisition of high-resolution spectral information beyond that of the visible spectrum. When applied to plants, it effectively enables non-invasive characterization of physiological status and has been widely used in agricultural settings. Marchantia is a model bryophyte species whose flat morphology and visually distinct stress-response phenotypes makes it an ideal candidate for imaging studies. Here, we provide a comprehensive protocol for hyperspectral imaging for Marchantia plants, which encompasses hardware configuration, data acquisition, and computations processing. This protocol features a streamlined data processing pipeline hosted on a web-based development platform that automates 1) the segmentation of plant area into spatially distinct regions for localized analysis of intra-specimen physiological gradients, and 2) classification of plant pixels based on their spectral signatures. All results are exported as structured CSV files for ease of further analysis as desired by the user.

## Introduction

Hyperspectral imaging (HSI) is a technique integrating spectroscopy and digital photography for simultaneous acquisition of spatially resolved spectral information^1,2^. Unlike the typical trichromatic photography, which perceives only the broad red, green, and blue bands on the visible light spectrum, HSI captures bands beyond the visible spectrum as hundreds of narrow contiguous bands of 5-10 nm ^2^. This enables the detection of unique spectral signatures associated with different chemical or biological properties; a feature which has been applied for non-invasive characterization of diverse matrices. Within plant science, HSI has been used extensively for monitoring of plant physiological status ^2,3^, namely detection of symptoms of abiotic stress, nutrient deficiency, and disease. Exhibiting a highly variable phenotype that is readily visualized due to its flat architecture, Marchantia presents an ideal subject for such imaging work.

Here, we present a protocol for the hyperspectral imaging of Marchantia covering the setup of the imaging platform, followed by data collection, processing, and analysis. Depending on the size and complexity of the setup, users may expect the initial setup to take at least an hour. Subsequent data collection is relatively rapid at approximately three minutes per sample. Data processing and analysis is done on a web browser interface via Google Colab, using a custom script that combines the functionalities developed in our previously published work ^3,4^. This custom script selects for and segregates leaf area into three radial regions, generates normalized spectral readings, and classifies plant pixels based on their spectral signatures. Normalised readings and classifications are exported as CSV files, which can be used for further analysis as required by the user.

## Materials

It is essential that you consult the appropriate Material Safety Data Sheets and your institution’s Environmental Health and Safety Office for proper handling of equipment and hazardous materials used in this protocol.

### Biological materials and reagents

- Marchantia plants
  - Plants can be grown under aseptic conditions or in the soil, but should ideally have a diameter of at least 0.5 cm. There should also be sufficient spacing between plants to avoid overlapping of thalli.

### Software

- Google Colab code (https://colab.research.google.com/drive/1_2oJu8bewhi1PlhVVRum3nC1oVZkKUPE?usp=sharing)
  - This code provides a user-friendly, web-based means of analysing hyperspectral data. The analysis will tap into several custom functions, which are uploaded into GitHub (https://github.com/dr-daisuke-urano/PlantHyperspectralSVD)
- Example images used in this protocol have previously been published by Krishnamoorthi et al. (2024) ^4^ and can be downloaded from https://github.com/dr-daisuke-urano/PlantHyperspectralSVD

### Equipment and supplies

- Room or space with ventilation and consistent lighting
  - While a dedicated cabinet for imaging is not necessary, consistent ambient lighting can help avoid introducing variability in captured reflectance, particularly if there are multiple experiments and/or timepoints. The imaging setup should therefore be located away from the influence of strong sources of variable light (e.g. direct sunlight). Halogen lamps also generate a lot of heat during operation, so proper ventilation is necessary.
- Stable work surface
- Hyperspectral camera
  - Model used will depend on the experimental requirements. Examples featured in this protocol were captured with a Specim IQ (CMOS sensor with 204 spectral bands across 400-1000 nm, image resolution 512 × 512 pixels) (Fig2A). However, our laboratory has also used the Specim FX10 (CMOS sensor with 224 spectral bands across 400-1000 nm, image resolution 1024 × 1024 pixels), and Specim FX17 (CMOS sensor with 224 spectral bands across 900-1700 nm, image resolution 640 × 640 pixels) for other work (Fig2B).
- Appropriate stand with camera mount allowing for top-down capture
  - A copystand allows for reproducible imaging at variable heights, which is ideal for capturing the flat morphology of Marchatia thalli. A tripod or other type of mount that can support the weight of the hyperspectral camera and allows for top-down angling can be used as a substitute, though care must be taken to reproduce the setup as exactly as possible between images and timepoints. Examples featured in this protocol were captured with a Specim IQ mounted on a copy stand (Fig2A).
  - A specialised setup may also be used if available. (e.g. Specim LabScanner 40×20 setup, on which the Specim FX10 and FX17 are mounted (Fig2B)).
- White reference tile
- Dark reference
- Plant growth vessel
- Matte, relatively non-reflective white surface with a solid colour (e.g. white printing paper)
- Halogen bulbs
- Appropriate mounts for halogen bulbs
- Appropriate stands for mounted halogen bulbs
  - Stands should be stable
- Extension cords, as required
- Rulers
- Computer with internet connection and web browser

**Figure 1.**
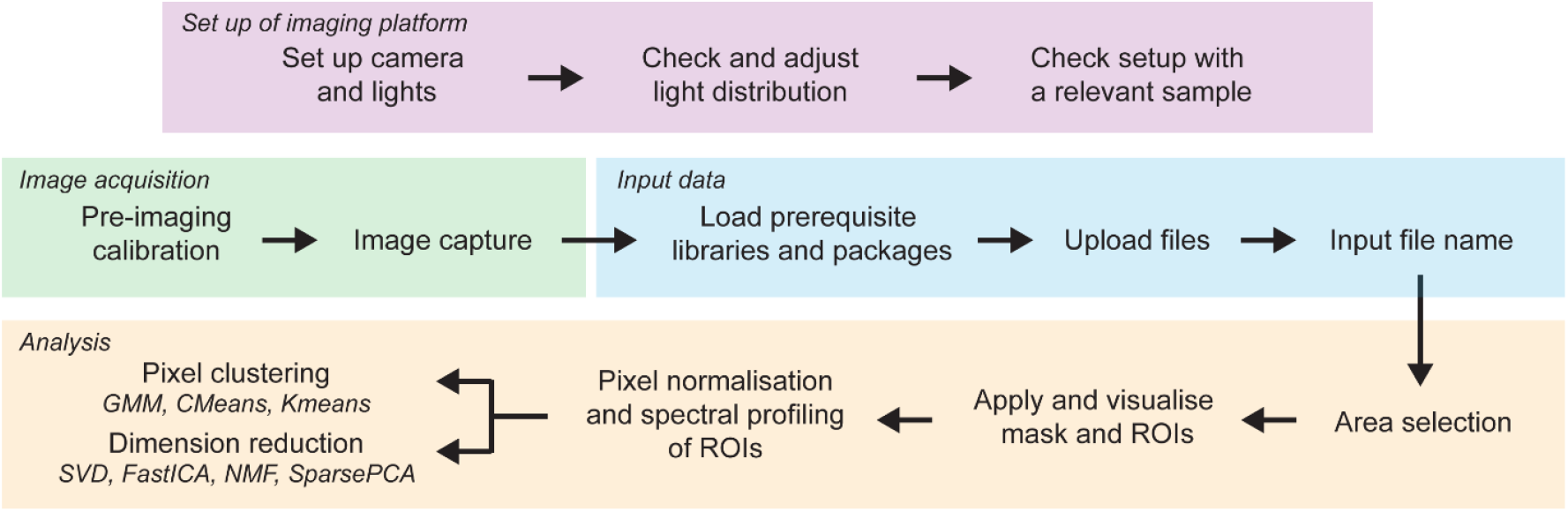
Overview of protocol from set up of imaging platform to image acquisition and data analysis.

**Figure 2.**
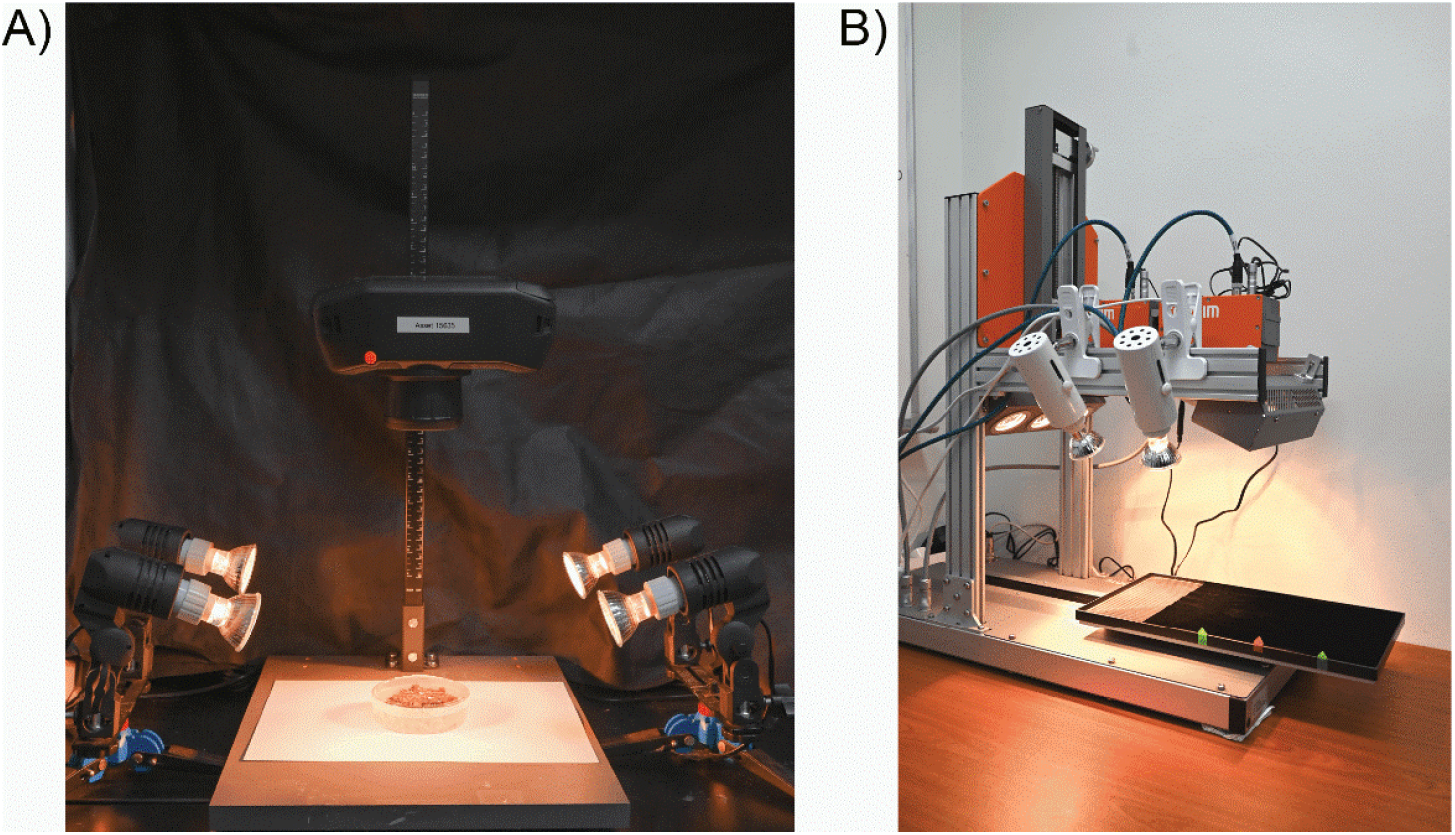
Hypersepctral imaging setups. A) Specim IQ camera mounted on a copystand with light provided by four lamps. B) Commercial Specim LabScanner 40×20 setup with inbuilt lighting and Specim FX10 and FX17 cameras mounted. Supplemental lighting was added in the form of two lamps clipped to the top of the setup.

## Method

### Set up of imaging platform

This setup process covers the manual mounting and positioning of the hyperspectral camera and light sources. Some commercial hyperspectral imaging setups come with camera and lamps preinstalled; however, light intensity and distribution should still be checked with in relation to the intended height and position of the actual sample (Step 5). If inadequate, supplementary lighting may be installed (Fig2B). Lighting is a significant source of variability and care should be taken to ensure the imaging area is as evenly lit as possible. Even with the best lighting however, minor variations may exist within the space – these can be corrected for during postprocessing.

1. Set up copystand on a stable surface, away from strong sources of variable light.
2. Mount hyperspectral camera on a copystand and turn the camera on. Set up the camera following the manufacturer’s instructions.
3. Mount halogen bulbs and place them around the copystand. Turn lamps on.
4. Using the camera’s viewfinder and a dummy sample or vessel, adjust camera height and focus, and mark out the ideal sample position on the copystand platform.
  a. Ensure that entirety of sample is visible. If sample is grown in a vessel (e.g. Petri dish, multi-well plate), check that the view is unimpeded by vessel walls. Multi-well plates for instance, may require multiple images at different positions to fully capture all samples.
5. Adjust positions of lamps such that illumination is relatively even across the sample position.
  a. Warning: Halogen lamps heat up rapidly and can pose a burn risk. Avoid direct contact with lamp during and for at least 10 mins after operation.
  b. Initial estimation of light distribution can be done visually with the aid of a non-reflective flat surface of consistent colour (e.g. white backdrop), or a dummy sample or vessel. More precise checks can be done by checking the variance in spectral readings at different spots within the image.
  c. Number and positions of lamps depend on the size of the sample and properties of individual bulbs. Smaller bulbs with lower power will naturally have a lower intensity and smaller illumination range; to compensate, more bulbs may be positioned around the imaging site at a closer distance. Lamps should ideally be angled downwards at about 40 to 80° relative to the base of the copystand, though smaller angles (down to 30°) can be used with flatter vessels like petri dishes and multi-well plates. In our setup, a total of four small halogen lamps (20W each) were positioned at the sides of the copy stand, two per side. Lamps on each side were approximately 20 cm apart, and angled inwards and downwards towards the imaging site (Fig2A).
  d. In arranging lamp positions, please also allow room for the operator to safely move and manipulate samples as needed.

### Optimisation of integration time

Most cameras can automatically determine optimal integration time. Depending on the samples, however, further adjustments may be required. Integration time should be adjusted to avoid over- and underexposure of samples, as that will distort reflectance values and affect accuracy of readings. A dummy sample with a similar phenotype to what is expected is recommended for this optimisation step.

6. Take an image of a dummy sample and check spectral readings at multiple points across the sample. Adjust integration time as required.
  a. Dark spots or regions produced by heavy pigmentation (such as during phosphate deficiency) and necrosis tend to be underexposed and will require higher integration times.

### Image acquisition

Once setup is confirmed and camera calibrated, refrain from altering the setup or lighting. Any disruption (e.g. lights being moved), will require re-calibration. Note also that hyperspectral imaging can detect water content, so ensure that plants are properly watered prior to imaging.

7. Turn on lamps and allow them to warm up for at least 5 minutes before calibration, or until light intensity has stabilised.
8. Perform calibration as required following the manufacturer’s protocol, with the white reference placed at the same plane as the sample.
9. Replace white reference tile with the sample. Adjust focus as required and proceed with imaging.
  a. Refrain from moving the sample during imaging, and do not block the lights.
  b. Halogen bulbs generate a lot of heat. To avoid unnecessary stress, plants should not be exposed to halogen lamps beyond the time required for imaging.

### Data extraction and analysis

The following steps detail the web-based processing and analysis of raw hyperspectral data using a Google Colab script. The following tasks are performed: selection and segregation of regions of interest (ROI), background masking, data normalization, pixel classification, and dimension reduction. ROI selection identifies plant area and subsets selected area into concentric circles for localized analysis. Masking isolates ROIs from the background based on a user defined threshold for further analysis. Data normalization serves to correct for non-biological variations that present even in the face of even lighting, such as uneven sample topography and hardware limitations (e.g. lens vignetting). Pixel classification groups pixels based on their reflectance spectrum. Dimension reduction simplifies the datacube for extraction of key features.

10. [Under Section 1] Run Cell 1 to install necessary libraries and import functions from Github
11. [Under Section 2] Run Cell 2 and upload Specim IQ data files. The following files are required
  a. img_id.hdr
  b. img_id.raw
  c. WHITEREF_img_id.hdr
  d. WHITEREF_img_id.raw
  e. DARKREF_img_id.hdr
  f. DARKREF_img_id.raw
12. [Under Section 2] In Cell 3, define image ID in `IMG_ID` and run the cell.
13. [Under Section 3] Run Cell 4 and select plants by clicking any pixel on the plants. (Fig 3A)
14. [Under Section 4] In Cell 5, define the threshold value for masking in `threshold_val` and run the cell.
  a. Check that the plant area is properly selected, with minimal inclusion of non-plant pixels (Fig 3B)
  b. A threshold value of 1.8 is sufficient for the example images, but may not necessarily work with user-supplied images. Users should adjust the values as needed.
15. [Under Section 5] Run Cell 6 to perform data normalisation and spatially-resolved spectral profiling (Fig 4).
  a. Data is normalized to the near-infrared (nIR) bands (875-925nm, indexed as 170-175 in dataframe)
  b. Extracted data is also divided into three concentric ROIs (“central”, “paracentral”, and “peripheral”) to generate localized spatial information.
  c. Extracted data is compiled into a .csv file.
16. [Under Section 6] Run Cell 7 to perform whole plant pixel clustering and spectral component analysis. Parameters for clustering and dimension reduction can be specified in their respective lines.
  a. For clustering, the number of final clusters and algorithm of choice can be specified under the `num_clusters` and `method` parameters, respectively. Three clustering methods are available: Gaussian mixture model (“GMM”), c-means (“Cmeans”), and k-means (“Kmeans”) (Fig 5A).
  b. Similarly for dimension reduction, number of dimensions and method of choice can be specified under the `dim` and `method` parameters, respectively. Four dimension reduction methods are available: Singular Value Decomposition (“SVD”), Fast Independent Component Analysis (“FastICA”), Non-negative Matrix Factorisation (“NMF”), Sparse Principal Component Analysis (“SparsePCA”) (Fig 5B).
17. [Under Section 7] Run Cell 8 to export all results as a ZIP folder.

**Figure 3.**
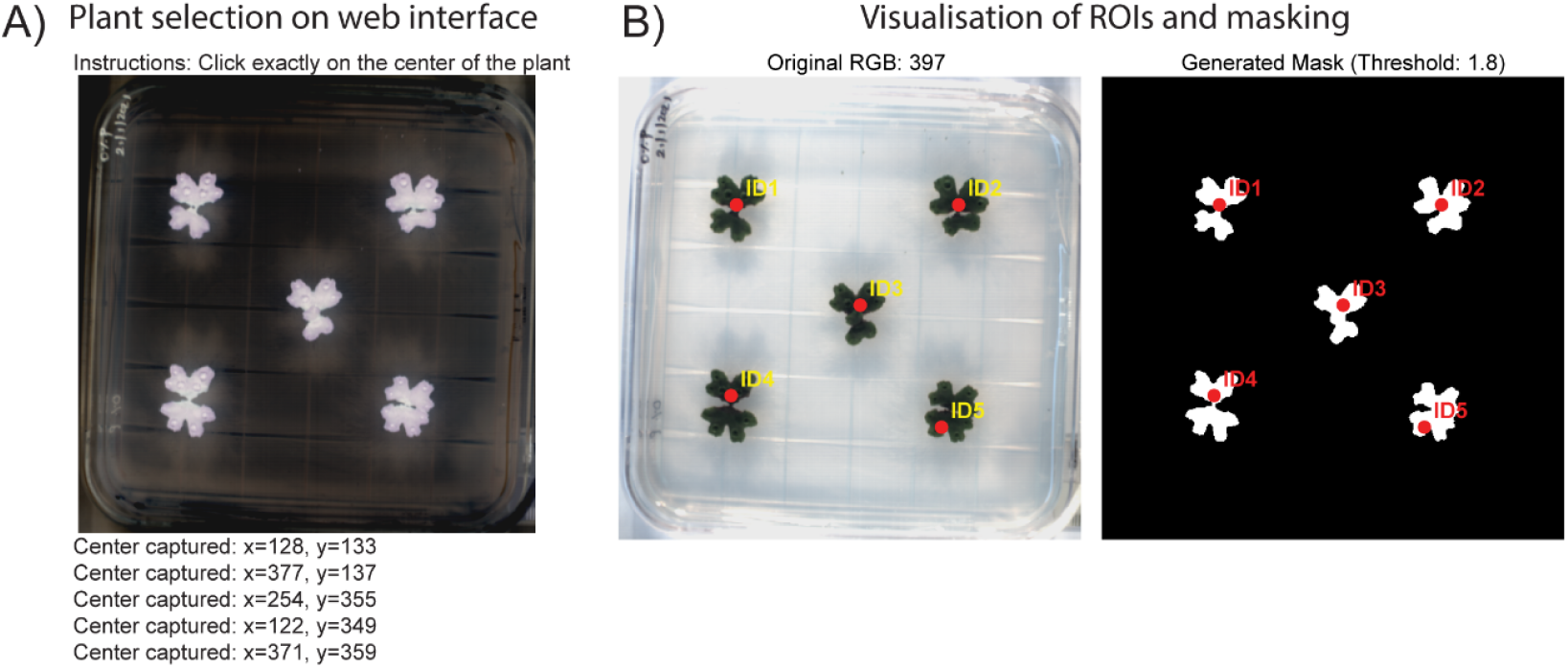
Example of web interface for A) plant selection and B) visualisation of masking results.

**Figure 4.**
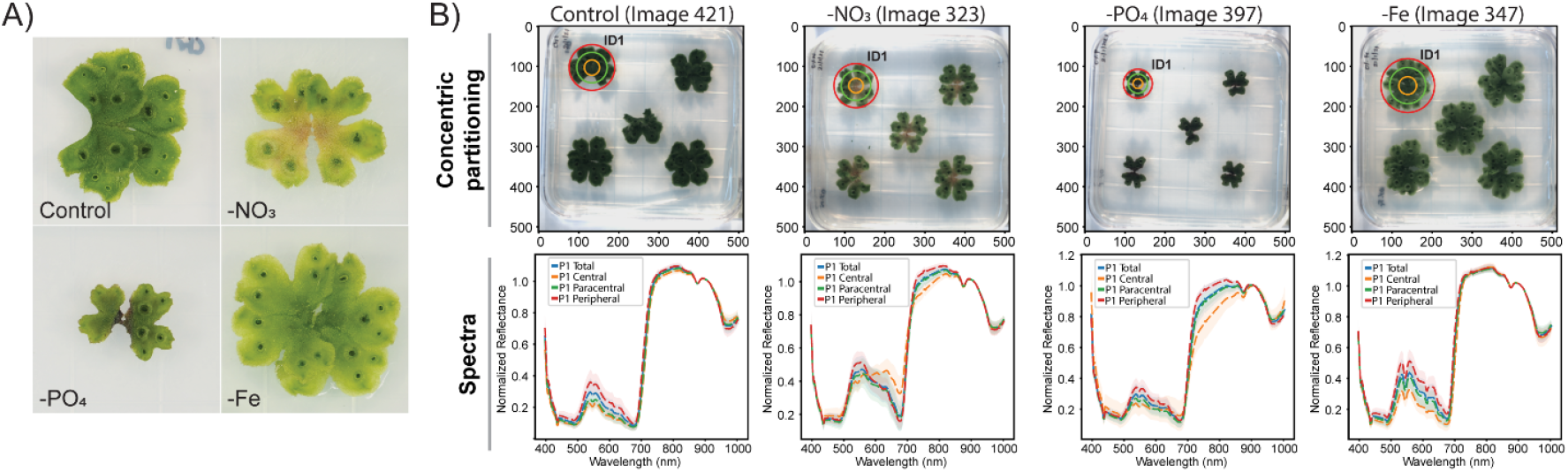
Representative samples and their spectral signatures. A) RGB images showing representative phenotypes of nutrient-stressed Marchantia. B) Extracted spectral signatures from a representative plant from each plate. Plant area is divided into three concentric ROIs: “central”, “paracentral”, and “peripheral”, indicated by the orange, green, and red lines respectively. Mean of extracted spectra from each ROI are presented in the line graphs following the same colour scheme, with mean of the whole plant ROI presented in blue. Lines represent mean values of pixels from the respective ROIs, while shaded areas represent standard deviation.

**Figure 5.**
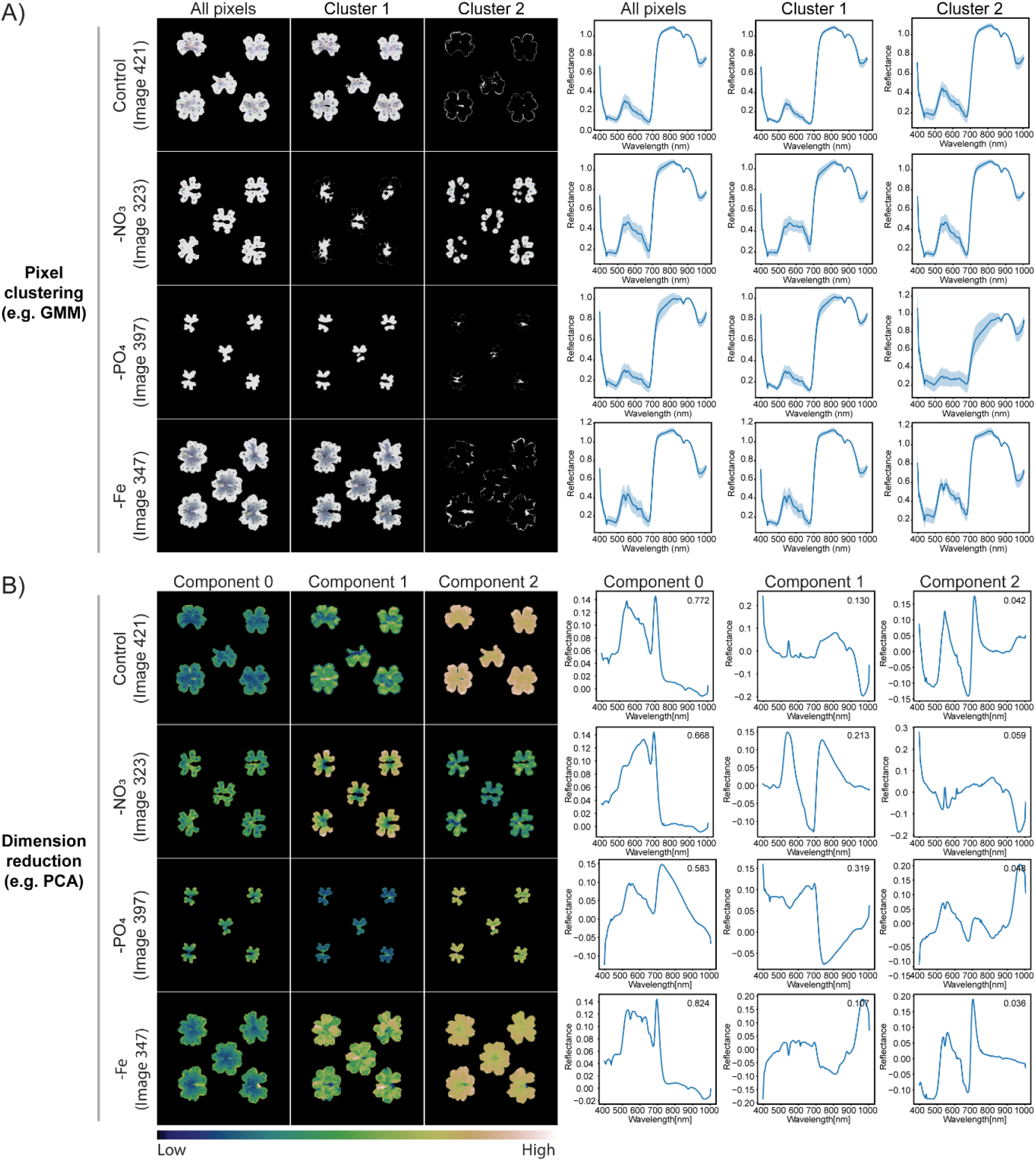
Example results demonstrating pixel clustering and dimension reduction. A) Pixel clustering using the GMM method. Grayscale images show the original image and the generated clusters, while plots show the respective spectral signals. Lines represent mean values of pixels from five plants, while shaded areas represent standard deviation. B) Dimension reduction via PCA. False-colour images show pixel intensity of selected ROIs from each cluster, while plots show the respective spectral signals. Lines represent mean in plot areas indicate the fraction of variance within the component.

## Troubleshooting

Problem (Steps 5): Uneven light intensities between lamps

- Solution: Check the bulb. Wear and tear will naturally lead to loss of light intensity overtime, and some bulbs may degrade faster, especially if bulbs are already of different ages. If light intensity becomes too low, replace the bulb. The new bulb will very likely have a higher light intensity compared to the others; this difference can be compensated for by adjusting the lamp placement and distance between lamp and sample. Problem (Step 9): Condensation within tissue culture vessel obstructing view
- Solution: Avoid sudden, large temperature differences when transporting plants to imaging setup, or placing plates directly under air conditioning vents. Refrain also from stacking plates before imaging. Should condensation persist, plates may be opened under aseptic conditions and the condensation blotted from the lids with a clean paper towel sprayed with 100% ethanol. Ensure lids are completely dry before covering the plate. Problem (Step 9): Plants drying out
- Ensure plants are sufficiently watered and placed away from the heat of the halogen lamps before and after imaging. If issues arise that disrupt the imaging process, remove the plant from the setup and return it only when ready for imaging. Problem (Step 9): Poor resolution of images
- Check that focus of the camera and adjust accordingly until samples are in focus. Check also that the distance between the camera and sample is outside the minimum operating distance of the camera. Problem (Step 12): FileNotFoundError message
- Check that value input into `IMG_ID` matches ID of uploaded files. Problem (Step 13): Plant not selected
- Ensure that plant pixel is clicked when selected ROI. Problem (Step 14): Plant area not properly selected
- Adjust threshold value. Problem (Step 16): Patterns presented in the cluster and/or component plots are indistinct or poorly matched to the apparent the visual pattern from RGB images
- Adjust number of groups and/or clustering method.

## Discussion

We previously published a hyperspectral imaging protocol for analysis of stress-induced colour changes in Marchantia ^4^, which was subsequently expanded to include additional data classification methods for pigmentation pattern analysis in ornamental plants ^3^. The analysis pipeline presented in this protocol is an amalgamation of their unique functionalities (i.e. data normalisation, ROI segregation, and pixel classification) into a user-friendly web interface. This is not only suitable for users with no programming background, but also circumvents the complications arising from different package versions. Offline analysis may still be done by downloading the original script and custom functions, which can be customised by experienced users to match their needs. Examples of results using previously published sample data ^4^ are illustrated in Fig 5.

Users should note that several limitations apply. Firstly, scripts and functions are adapted for files generated with SPECIM hyperspectral cameras and may not work with files generated from other camera models with different naming conventions or formats. Secondly, this pipeline is best utilized on images captured against a solid white background, as the masking and ROI selection functions are based on contrast in the blue channel of the RGB image. Use of other background colours may not provide the same level of contrast. Deviation from any of the above would require manual adjustment of the script and functions for proper file recognition and plant area selection.

While not necessarily applicable to all circumstances, this pipeline nonetheless embodies the core steps in hyperspectral data analysis: 1) data normalization, 2) ROI selection, and 3) data classification. The specific methods are by no means prescriptive as analysis is heavily context-driven, and can be adjusted according to the research question.

Several methods exist for data normalization, which serves to reduce noise from non-biological influences which can affect downstream classification (Fig 6). Here, pixel normalization was performed using the mean reflectance from the 875-925 nm wavelengths as it exhibited the lowest variance across samples and the generally high signals across species and stresses ^4^. Following normalization, background masking is performed to remove extraneous pixels in preparation for classification. Selected ROIs can also be further divided for more detailed spatial information. For Marchantia, concentric segregation was employed as symptom development tended to occur in a radial pattern from the central part of the plant under the conditions tested (i.e. nutrient stress) ^4^. Subsequent classification can be performed on the subsetted ROIs, or on the whole plant (example in Fig 6), depending on the research question. The specific classification and dimension reduction methods to be used are dependent on the data, and should be determined by the user. Descriptions of these methods can be found in Krishnamoorthi and Urano (2025).

**Figure 6.**
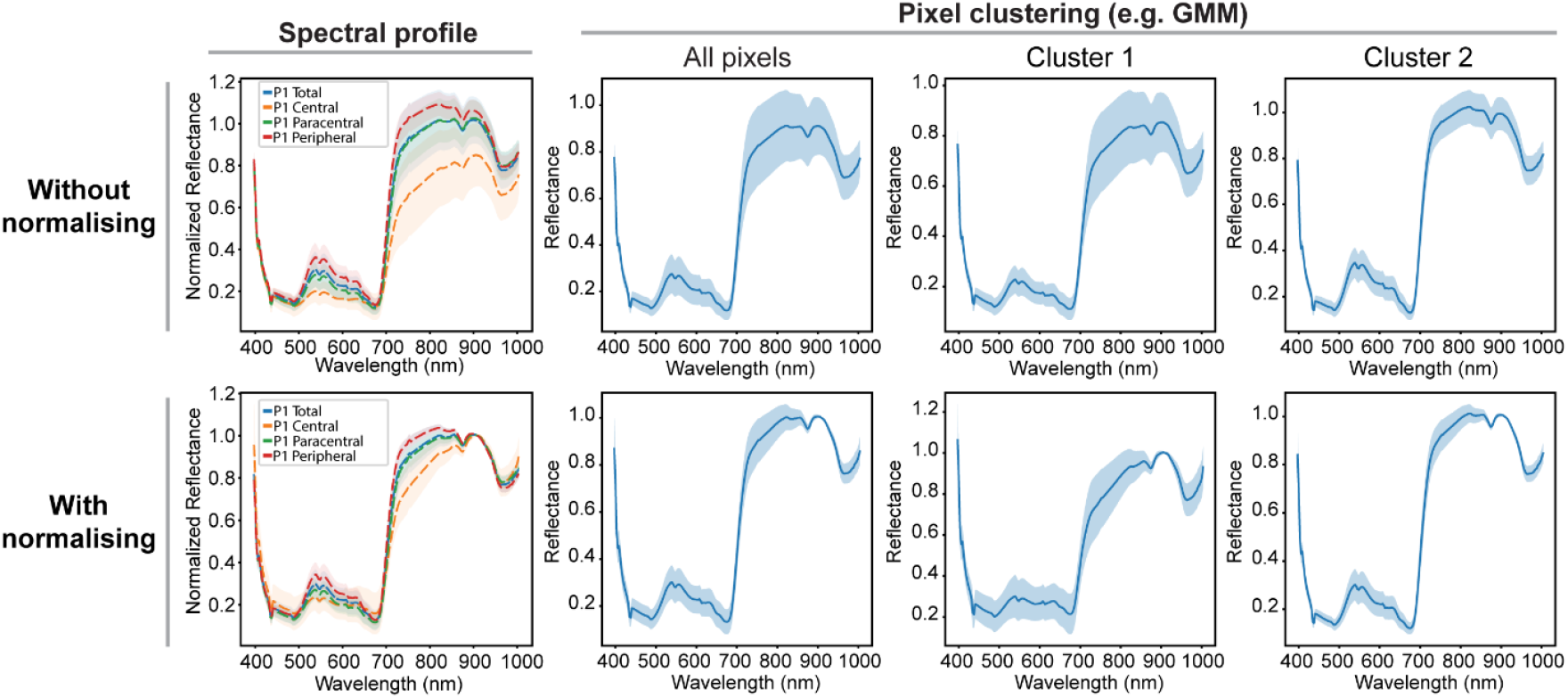
Impact of pixel normalization on the spectral profile and downstream clustering. The example spectral profile shown here is from the representative plant under phosphate deficiency in Fig 4. Dotted lines represent mean of pixels from the ROIs, while shaded areas represent standard deviation. The example used for pixel clustering by GMM shown here is from the representative phosphate deficiency treatment in Fig 5. Solid lines represent mean of pixels from the five plants, while shaded areas represent standard deviation.

## Competing interest statement

The authors declare no competing interests.

## Acknowledgements

The authors would like to thank Ms Krishnamoorthi Shalini for the sample images used in the examples featured in this protocol. This study was supported by the Agency for Science, Technology and Research (A*STAR) Singapore under the industry alignment fund pre-positioning program; High Performance Precision Agriculture system (A19E4a0101), and by the National Research Foundation (NRF) Singapore under its Campus for Research Excellence and Technological Enterprise (CREATE) program. Both grants were granted to DU.

## Author contributions (CRedIT)

Conceptualisation: DU. Methodology: DU, GZHT. Software and data curation: DU. Writing – original draft: GZHT. Writing – Review and editing: GZHT, DU. Visualization: GZHT. Supervision, project administration, and funding acquisition: DU

## Notes

### Competing Interest Statement

The authors have declared no competing interest.

